# A new, phenotypically distinct subpopulation of Regulatory Killer T ex-Th17 cells expressing CD4^low^CD25^hi^CD49^hi^Foxp3^hi^ROR^low^IL-17^low^

**DOI:** 10.1101/261073

**Authors:** Maria S. Sayapina

## Abstract

Th17 and regulatory T (Treg) cells are integral in maintaining immune homeostasis and Th17– Treg imbalance is associated with inflammatory immunosuppression in cancer. Here it is shown that in addition to ROR+Foxp3+ cells eTreg cells are a source of ex-Th17 CD4^low^CD25^hi^CD49^hi^Foxp3^hi^ (Regulatory Killer T – RKT) cells while the latest are much more suppressive. Moreover, we have identified a set of key cytokines that favor the generation and expansion ex-Th17 Foxp3^low^ cells. These findings should accelerate efforts to define the function of this new subset of Treg cells in the immune response to cancer.

## Introduction

Treg cells consist of functionally diverse subsets of immune suppressive T cells that play a crucial role in the modulation of immune responses and the reduction of deleterious immune activation [1, 2]. Treg cells may participate in the progression of cancer, especially with regard to the ability of Treg cells to promote the development of tumors [3]. It was described that the levels of intratumoral Treg cells correlating with better or worse outcomes depending on the tumor type [4, 5]. Recent studies indicate that human ovarian cancer cell line SKOV-3 could convert, in the presence of IL-2, Treg into Th17 cells. These results support the ability of the tumor microenvironment to regulate and expand IL-17-producing T-helper (Th17) cells. Similar results were obtained upon stimulation of CD4+ T cells in the absence of tumors but in the presence of IL-1β/IL-6 and IL-2 [6]. Cytokine profile analysis revealed that ovarian tumor cells, tumor-derived fibroblasts, and antigenpresenting cells (APCs) secrete IL-1β/IL-6 [7]. IL-1β is a potent inducer of Th17 cell differentiation and expansion, whereas IL-6 is capable of expanding memory Th17 cells [8]. Gene profile analysis revealed that SMAD 6 and HDAC 11 are hyperexpressed in ovarian cancer cell line SKOV-3 [9]. In its turn Smad6 is transcriptionally induced by the antiinflammatory cytokine TGF-β. On the one hand, the importance of the concentration of TGF-b was illustrated in selectively regulating Treg and Th17development. Low concentrations of TGF-B favor Th17 differentiation by enhancing IL- 23 receptor (IL- 23 R) expression, while high concentrations promote Treg differentiation by inhibit IL-23 R up-regulation [10]. On the other hand, it was also described that ovarian tumor cells secreted a high amount of latent TGF-β (inactive form), but the level of active form of TGF-β was very low (≤30 pg/ml) or undetectable because of its short half-life [8, 11]. Importantly, most of all types of tumors secrete high amount of TGF-β. Whereas overexpression of HDAC11 was associated with inhibition of expression of the gene encoding IL-10 and higher IL-12 mRNA expression [12]. IL-12 and IL-23 shared the same IL-12Rβ1 receptor subunit and are characterized by overlapping effects on target cells. As shown before, IL-23 stimulation is not only crucial for attaining full effector function but also necessary for double expression of IL-17A and IFN-γ, induction of T-bet and subsequent deviation toward IFN-γ production[13]. Here, it was provided an insightful mechanism by which ex-Th17 CD4^low^CD25^hi^CD49^hi^Foxp3^hi^ cells are generated from eTreg cells and regulated by cytokines that favor the generation and expansion exTh17Foxp3^low^ cells.

## Results

I conducted 3 experiments aimed at deriving eTreg cells using BALB/c mice (1^st^ time) and Foxp3-GFP-DTR (2 times), CD4+CD25+FOXP3^DTR-GFP^ cells were isolated from lymph nodes and spleen by flow cytometry cell sorting to high purity and stimulated with anti-CD3/CD28 coated Dynobeads and IL-2 (Fig. 1). FACS analyses of the isolated population was at day 3 after stimulation. As shown in Figure 2, ex-Th17 CD4^low^CD25^hi^CD49^hi^Foxp3^hi^ cells were clearly detectable in populations from the purified CD4+CD25+ T-cell fractions after in vitro expansion. Staining with anti-IL-17 antibody revealed that ex-Th17 CD4^low^CD25^hi^CD49^hi^Foxp3^hi^ cells secreted low level of IL-17, although ROR+FOXP3+ T cells produced high level of IL-17 Further experiments revealed that freshly isolated CD4^low^CD25^hi^T cells were strongly positive for CD49b and Foxp3 molecules and weakly positive for ROR (Fig. 3).

**Figure 1.**
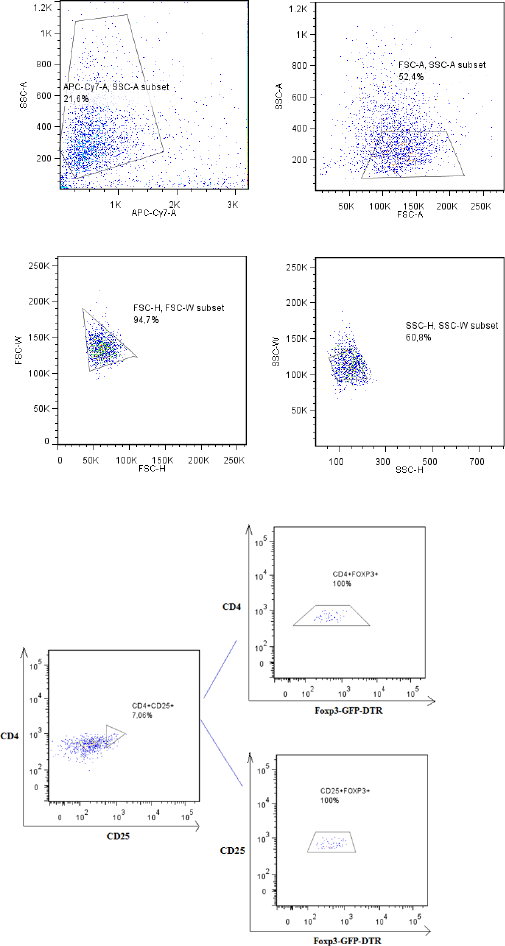
Sorting strategy and the purities of resultant population CD4+CD25+Foxp3+T cells.

**Figure 2.**
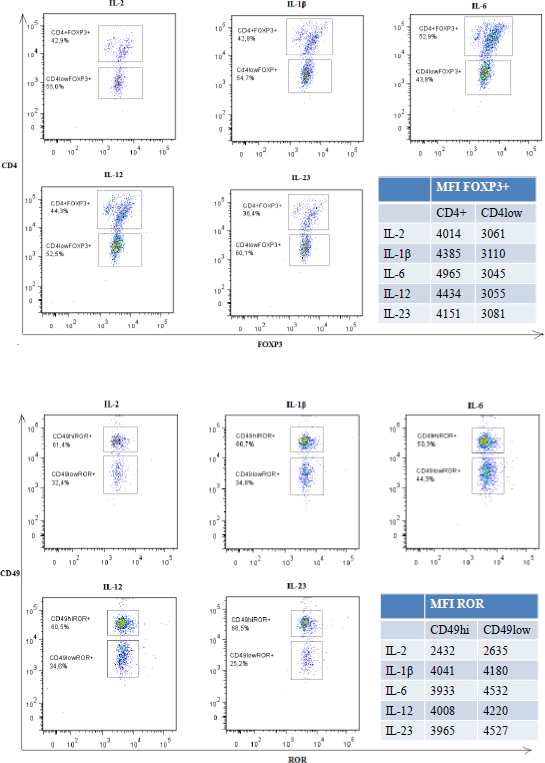
CD4^low^ T cells express Foxp3 and CD49b.

So I am the first who describe this subpopulation of ex-Th17 CD4^low^CD25^hi^CD49^hi^Foxp3^hi^ cells.

**Figure 3.**
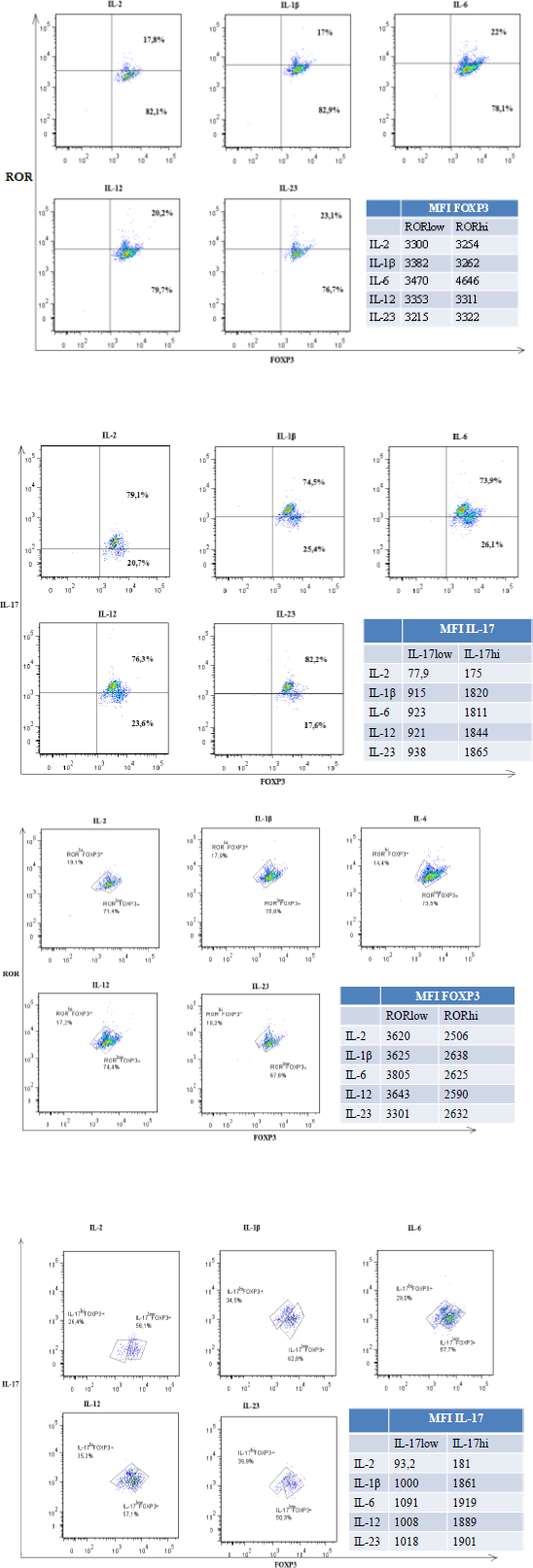
CD4^low^CD25^hi^CD49^hi^Foxp3^hi^ cells express low level of IL-17 and ROR.

We also sought to determine the role of key cytokines, as discussed earlier, in the generation and expansion of exTh17Foxp3^low^ cells.

In the next stage we tested the effects of IL-12, IL-1β, IL-6, IL-23 on exTh17Foxp3^low^ differentiation and expansion by using Treg cells from Foxp3^DTR^-GFP mouse. IL-2-containing medium provided a baseline for comparison. Analysis of ex-Th17 CD4^low^CD25^hi^CD49^hi^Foxp3^hi^ cells revealed that IL-23 plays a more prominent role in differentiation and expansion of exTh-17 Foxp3^low^ cells than do IL-12 and IL-1β. By contrast, IL-6 stimulated IL-17-producing ROR+Foxp3+T suppressive cells.

## Discussion

In this work, we have described the new subpopulation of ex-Th17 CD4^low^CD25^hi^CD49^hi^Foxp3^hi^ cells. Importantly, our data demonstrate the differentiation of eTreg cells to ROR+Foxp3+ cells and exTh17 RKT cells. Bryl et al. have previously reported population of peripheral blood T cells with reduced CD4 and high CD25 expression (CD4^low^CD25^high^), that are able to non-specifically suppress the proliferation of autologous, previously polyclonally activated CD4+lymphocytes and to kill them by direct contact. CD4^low^CD25^high^ T cells expressed significant amounts of both intracellular perforin and granzyme B. At the same time common NK/NKT antigens, including: CD16, CD56, CD94, CD158b, CD161 and invariant NKT (iNKT), - were not present on CD4^low^ T cells [14].

Also using whole-genome microarray data sets of the Immunological Genome Project, it was demonstrated a closer transcriptional relationship between NK cells and T cells than between any other leukocytes, distinguished by their shared expression of genes encoding molecules with similar signaling functions, including NT cells and Treg [15]. In terms of common expression of Zap70 and Prscq and potential expression of perforin and granzyme B we concluded that the definition of a CD4^low^CD25^hi^CD49^hi^Foxp3^hi^ cells phenotype is enough to unambiguously detect and study the regulatory function of new subpopulation called Regulatory Killer T – RKT cells which fulfils the current phenotypic criteria identifying the exTh17 RKT cells by simultaneously expressing low amounts of ROR and IL-17A.

Thus, the relative concentration of IL-2, IL-12, IL-1β and IL-23 in the tumor microenvironment may be a critical factor for the generation of exTh17 RKT that will be converted into INF-γ- producing exTh17Foxp3^low^ (exTh17/Th1) cells. Based on these findings, it was predicted that cytokine milieu (low amounts of TGF-β and high amounts of IL-2, IL-12, IL-1β and IL-23) in cancer favors the generation and expansion of exTh17Foxp3^low^ cells, although further studies are needed to validate this concept.

In terms of several types of tumors secrete some cytokines, for example colorectal cancer express high level of IL-23, ovarian cancer – IL-12 (I am planning to prove it) the combination of IL-2, IL-12, IL-1β and IL-23 in different ways enhanced the differentiation of exTh17Foxp3^low^ (exTh17/Th1) cells from eTreg cells while retaining their ability to expand ROR+Foxp3+ T cells. IL-23 as a critical factor driving exTh17Foxp3^low^ cell expansion. Our findings clearly support the emerging concept that tumor environmental factors drive the generation and expansion of exTh17Foxp3^low^ cells. This knowledge should accelerate efforts to describe the new subpopulation ex-Th17 CD4^low^CD25^hi^CD49^hi^Foxp3^hi^ (RKT) cells in more detail and create several drugs for several immunogenic types of tumors (melanoma, ovarian cancer, renal cancer, colorectal cancer) on the basis of IL-2, IL-12, IL-1β and IL-23 that will be delivered locoregionally (intraperitioneally, intrahepatic artery etc) to decrease systemic toxicity.

## Methods

### Cell culture

CD4+Tcells that were isolated from FDG mouse by negative selection with mouse CD4+Isolation Kit and were further separated into CD4+CD25+FOXP3^DTR-GFP^+eTregcells using a FACS ARIA II instrument. Sorted 1,2×10^5 eTreg cells were cultured in the presence of anti-CD3- and anti-CD28-coated (2,5 mcl) Dynabeads and IL-2 (100U/ml). In some cultures IL-12 (30ng/l), IL-1β (30ng/ml), IL-6 (30ng/ml), IL-23 (30 ng/ml) were added. Cells were analysed with an FACS Canto instrument 3 days later.

Cells were cultured in culture medium (RPMI-1640 supplemented with 100 U/mL penicillin, 100 g/mL streptomycin, 5 mM 2-mercaptoethanol, 0.05% and 10% fetal bovie serum [FBS]) at 37°C, and 5% CO2, in 96-well round-bottom plates (Greiner, Frickenhausen, Germany).

### Antibodies and reagents

Allophycacocyanin (APC)- and Cy7- conjugated anti-CD4 (RM 4-5) mAb, phycoerythrin (PE) – and Cy7-conjugated ant-CD4(RM 4-5) mAb, peridinin chlorophyll protein complex (PerCP)- andCy5.5-conjugated anti-CD25 (PC-61) mAb, phycoerythrin (PE) anti-ROR (Q31-378) mAb were purchased from Bioscience. Brilliant Violet 421(BV421) – conjugated anti-IL-17 (TC-11-18H 10.1) mAb, Alexa Fluor 647-conjugated anti-CD49b (DX5), LIVE/DEAD Fixable Near-IR Dead Cell Stain Kit were purchased from Biolegend. Recombinant murine IL-6, IL-12, IL-23, IL-1β were purchased from Sarstedt and Bioscience.

All results were received by the author personally in 01.2018 and reported on a scientific club in the laboratory on January 29, 2013, and this was the continuation of the author’s dissertation work [16].

## Acknowledgments

I thank Professor Shimon Sakaguchi for his encouragement and support.

